# Interaction between photoperiod and variation in circadian rhythms in tomato

**DOI:** 10.1101/2022.01.14.476322

**Authors:** Yanli Xiang, Thomas Sapir, Pauline Rouillard, Marina Ferrand, José M Jiménez-Gómez

## Abstract

Many biological processes follow circadian rhythmicity and are controlled by the circadian clock. Predictable environmental changes such as seasonal variation in photoperiod can modulate circadian rhythms, allowing organisms to adjust to the time of the year. Modification of circadian clocks is especially relevant in crops to enhance their cultivability in specific regions by changing their sensibility to photoperiod. In tomato, the appearance of mutations in *EMPFINDLICHER IM DUNKELROTEN LICHT 1* (*EID1*, Solyc09g075080) and *NIGHT LIGHT-INDUCIBLE AND CLOCK-REGULATED GENE 2* (*LNK2*, Solyc01g068560) during domestication delayed its circadian rhythms, and allowed its expansion outside its equatorial origin. Here we study how variation in circadian rhythms in tomato affects its perception of photoperiod. To do this, we create near isogenic lines carrying combinations of wild alleles of *EID1* and *LNK2* and perform transcriptomic profiling under two different photoperiods. We observe that EID1, but not LNK2, has a large effect on the tomato transcriptome and its response to photoperiod. This large effect of EID1 is likely a consequence of the global phase shift elicited by this gene in tomato’s circadian rhythms.

## Introduction

Synchronization with the environment is crucial for survival. In order to efficiently anticipate predictable environmental changes linked to diurnal and seasonal oscillations, all living organisms have developed endogenous timekeeping mechanisms named circadian clocks (Edgar et al., 2012). In plants, the circadian clock ensures the correct timing of crucial biological processes, such as growth, development, reproduction, photosynthesis or defense, just to name a few (Steed et al., 2021).

The circadian clock in plants is best studied in Arabidopsis, where it is organized in inter-connected regulatory loops (Harmer, 2009). In this species, two important transcription factor families are at the core of the clock. The REVEILLE (RVE) family, a set of MYB-like proteins that includes CCA1/LHY, RVE6, RVE3 and RVE8 (Rawat et al., 2011); and the PSEUDORESPONSE REGULATOR (PRR) family, composed of TOC1, PRR3, PRR5, PRR7 and PRR9 that are sequentially expressed during the day (Nakamichi et al., 2010). Additional members of the circadian clock include proteins that enable environmental perception such as ZEITLUPE, GIGANTEA, EARLY FLOWERING 3, EARLY FLOWERING 4, LUX ARRHYTHMO or the NIGHT LIGHT-INDUCIBLE AND CLOCK-REGULATED family (LNKs) (Sanchez et al., 2020). Most of these proteins are involved in light signaling and contribute to dynamically synchronize the circadian clock with the external cycles (Webb et al., 2019). For example, these proteins reset the circadian clock each day by reading light and temperature signals at dawn or dusk (Jones, 2009). Moreover, they can change the time of the day when some transcripts are expressed depending on the duration of the light period (photoperiod), allowing plants to time biological processes differently depending on the season (Dalchau et al., 2010; Michael et al., 2008; Miwa et al., 2006; Roden et al., 2002; Webb et al., 2019; Weng et al., 2019).

Because circadian rhythms can be modulated by the environment, their variation could be beneficial for adaptation to certain settings, and opens the possibility to selective pressures on the circadian clock. For example, plants that live near the equator where the duration of the day is constant along the year, could evolve different mechanisms to time their biological processes than plants at higher or lower latitudes that experience strong seasonal variation. Indeed, it has been found that plants adapted to higher latitudes naturally present longer free running circadian periods (Greenham et al., 2017; Hut et al., 2013; Michael et al., 2003). Moreover, allelic variation at circadian clock genes have been selected during crop domestication or breeding (Bendix et al., 2015; McClung, 2021). In soybean, variation in circadian clock genes such as GI (Watanabe et al., 2011), PRR3 (Li et al., 2019) or ELF3 (Lu et al., 2017) has been selected to control flowering time responses to photoperiod. The same has occurred in sugar beet with PRR3/7 (Pin et al., 2012), in barley with PPR3/7 (Turner et al., 2005) and ELF3 (Faure et al., 2012), and in pea and lentil with ELF3 (Weller et al., 2012).

Tomato is an interesting case because domestication significantly decelerated circadian rhythms through selection of knock-out alleles of two genes not identified as a source of variation in any other crop, *EID1* and *LNK2* (Müller et al., 2016, 2018). EID1 is an F-box protein that targets phytochrome A for degradation in Arabidopsis (Marrocco et al., 2006). In tomato, a three base pair deletion in *EID1* appeared early during domestication causing a ∼2 hour phase delay of its circadian rhythms (Müller et al., 2016, 2018). LNK2 is part of a family of light inducible coactivators in Arabidopsis that interact with proteins of the RVE family to regulate the expression of clock genes (Ma et al., 2018; Pérez-García et al., 2015; Rugnone et al., 2013; Xie et al., 2014). In cultivated tomato, *LNK2* presents a large deletion that increases the period of its circadian rhythms (Müller et al., 2018). The genomic regions of *EID1* and *LNK2* show signatures of positive selection and their mutations are fixed in cultivated tomatoes but do not exists in its closest wild ancestor *S. pimpinellifolium*, suggesting that phase and period changes have been beneficial for its domestication (Müller et al., 2016, 2018). Because both EID1 and LNK2 are involved in light signaling to the circadian clock, and tomato domestication started in the equatorial region of South America with constant photoperiods of ∼12 hours, it has been hypothesized that mutations in *EID1* and *LNK2* allowed tomato to expand its cultivation range to higher latitudes (Müller et al., 2016, 2018).

Here we study the interaction between photoperiod and the mutations that delayed circadian rhythms in tomato. For this, we generate a set of tomato near isogenic lines that segregate for wild functional alleles of *EID1* and *LNK2*. We first confirm the validity of these lines and then study their transcriptional responses to variation in photoperiod and to the allelic configurations of *EID1* and *LNK2*. Our study contributes to understanding how variation in circadian rhythms affects the molecular state of tomato under different photoperiods.

## Methods

### Plant Material

Tomato near isogenic lines LNK2+, EID1+ and LNK+/EID1+ (hereafter called LE+) were generated from a set of introgression lines derived from a cross between *S. lycopersicum* cv Moneymaker and *S. pimpinellifolium* accession TO-937, kindly provided by A. Monforte (Barrantes et al., 2014, 2016). Introgression lines SP1-2 and SP9-2 from this population, containing *S. pimpinellifolium* alleles in the region of *LNK2* and *EID1*, were backcrossed to the cultivated parent Moneymaker. Recombinants that reduced the introgressed regions were selected for two consecutive generations using markers detailed in Table S1, screening a total of 264 plants per line. Independent lines with reduced introgressions containing *S. pimpinellifolium* alleles of *EID1* and *LNK2* were crossed, and a whole genome sequencing was performed in a single F1 individual. Genotyping of short reads revealed the presence of introgressions in chromosomes 1 (the region of *LNK2*), 4, 5 (two introgressions) and 9 (the region of *EID1*) (Figure S1). The segregating progeny of this F1 line was genotyped for all these regions using markers detailed in Table S1 to select lines that had only the introgression in chromosome 1 containing *LNK2* (LNK2+), only the introgression in chromosome 9 containing *EID1* (EID1+) or introgressions only in chromosomes 1 and 9 (LNK2+/EID1+, hereafter LE+). In addition, all plants screened contained *S. pimpinellifolium* alleles on the distal region of chromosome 4 due to the presence of a segregation distortion locus identified during the generation of the population (Barrantes et al., 2014, 2016).

### Genotyping tomato lines

A single F1 individual resulting from the cross of two lines carrying reduced *S. pimpinellifolium* introgressions in the region of *LNK2* and *EID1* (see above) was sequenced for whole genome genotyping. To do this, DNA from leaf tissue was extracted using the DNeasy Plant extraction kit from Qiagen following manufacturer’s instructions. A single sequencing library was constructed using the standard Illumina method, and sequenced in an Illumina NovaSeq 6000 system, yielding 361,774,304 pairs of 150bp reads. In order to assign alleles in the F1 hybrid to cultivated and wild tomato we obtained publicly available short reads from *S. pimpinellifolium* accession LA1589 and *S. lycopersicum* cv. Moneymaker (Aflitos et al., 2014; Lin et al., 2014). Reads from all three genotypes were independently aligned to the tomato reference genome v2.5 using Bowtie2 version 2.3.4.2 with default parameters (Langmead and Salzberg, 2012). Previous to variant calling, reads with mapping quality lower than 5 were discarded using samtools v1.7 (Li et al., 2009), duplicated reads were removed using Picardtools (http://broadinstitute.github.io/picard) and indels were realigned using GATK IndelRealigner (McKenna et al., 2010). We called variants in all three alignments simultaneously using GATK UnifiedGenotyper (McKenna et al., 2010). Presence/absence of *S. pimpinellifolium* alleles in the F1 hybrid was scored at 3,396,011 biallelic positions where Moneymaker and LA1589 were homozygous and different from each other, and the hybrid was heterozygous.

### Leaf movement essays

Three different experiments to measure leaf movements were conducted following a protocol that has been described in detail elsewhere (Müller and Jiménez-Gómez, 2016). Briefly, seedlings were entrained in an environmental chamber to 12 hours light / 12 hours dark and 24 °C for seven to ten days. On the last day of entrainment, a polystyrene ball was attached to the tip of one cotyledon of each seedling and the conditions changed to constant light and temperature. Images of the seedlings were taken each 30 minutes for five days in constant conditions using Pentax Optio WG-1 digital cameras. Image analysis to extract the vertical position of the polystyrene ball was performed using ImageJ (Müller and Jiménez-Gómez, 2016; Schneider et al., 2012). Estimates for circadian variables were obtained via fast Fourier transform nonlinear least-squares analysis using BioDare2 (biodare2.ed.ac.uk) (Zielinski et al., 2014). Only seedlings with error (ERR) values below 0.4 were considered for statistical analyses.

### RNA-seq analysis

RNA-seq was conducted in two consecutive experiments in the same controlled environment chamber set first to LD (16 hours light / 8h dark, 24C) and then to SD (8 hours light / 16h light, 24 C). We collected leaf samples from 14-day old plants 2 hours after dawn (ZT2), coinciding with the maximum peak of expression of *EID1* and *LNK2*. Three biological replicates were collected from each genotype and condition, and total RNA was extracted with the RNeasy Plant Mini Kit from Qiagen. Libraries were prepared according to the Illumina TruSeq RNA protocol and sequenced in two lanes of a Novaseq 6000 system, yielding 834,954,102 150bp read pairs (per sample average of 34,789,754, minimum of 19,797,369).

Reads for each biological replicate were aligned independently to the tomato genome reference sequence v4.0 using hisat2 v2.1.0 (Kim et al., 2015), allowing a maximum intron length of 115400bp (the largest intron in the tomato genome annotation ITAG4.1). An average of 92,9% of the reads aligned to the reference (minimum of 89,8%). The number of reads per transcript in the ITAG4.1 annotation was counted with the featureCounts function in the Rsubread R package with default parameters (Liao et al., 2019). Only transcripts that presented more than 10 reads in all samples together were used in downstream analysis, leaving us with 23226 transcripts out of the 34688 transcripts present in the annotation. Sample homogeneity was surveyed the with the plotPCA function in the DEseq2 package in R (Love et al., 2014). Differential expression between each genotype and condition was calculated with DEseq2 using two different models. The first model aimed to obtain lists of differentially expressed transcripts between each genotype and *S. lycopersicum* cv Moneymaker in each photoperiod separately, and contained a unique variable with 8 factors grouping the genotype and condition for each sample (“MM_LD”, “MM_SD”,”LNK2_LD”, “LNK2_SD”, “EID1_LD”, “EID1_SD”, “LE_LD” and “LE_SD”). The second model was conceived to test the effect of photoperiod in each genotype and included the three variables “photoperiod”, “genotype at EID1” and “genotype at LNK2”, as well as all its possible interactions (photoperiod + EID1 + LNK2 + photoperiod:EID1 + photoperiod:LNK2 + EID1:LNK2 + photoperiod:EID1:LNK2). From this model we extracted the effect of photoperiod in each genotype. In both models we considered as differentially expressed those transcripts with a q-value lower than 0.01. A dataset with normalized values for each sample used in all downstream analyses and graphical representations was generated using the vst function in the DEseq2 R package (Love et al., 2014).

### Homologs of the PSEUDO-RESPONSE REGULATOR and REVEILLE gene families in tomato

To obtain the homologs of PRRs and RVE genes in tomato, we compared the protein sequences from the following Arabidopsis TAIR10 ids: AT1G01060.1 (LHY), AT1G18330.2 (RVE7), AT2G46830.1 (CCA1), AT3G09600.1, (RVE8), AT5G02840.1 (RVE4), AT5G17300.1 (RVE1), AT5G37260.1 (RVE2) AT5G52660.2 (RVE6), AT2G46790.1 (PRR9), AT4G18020.1 (APRR2), AT5G02810.1 (PRR7), AT5G24470.1 (PRR5), AT5G60100.2 (PRR3) and AT5G61380.1 (TOC1) onto the tomato protein sequences from annotation ITAG4.1 using standalone BLAST+ (Camacho et al., 2009). For the RVE family, we selected all matching tomato proteins with a bit score greater than 90, and for the PRR family, we selected those with a p value lower than 1e-40. Only tomato transcripts whose expression showed oscillation during the diel cycle were considered homologs. Retrieval and management of sequences was performed with the seqinr package in R [17] and neighbor-joining phylogenetic trees were constructed and drawn with the ape package (Paradis and Schliep, 2019) and the ggtree package (Yu et al., 2016) in R respectively.

### Determination of cycling genes in tomato

Short reads from a time course experiment performed in 7-day old *S. lycopersicum* var. M82 and *S. pennellii* seedlings were obtained from the SRA database (www.ncbi.nlm.nih.gov/sra, project number PRJNA295848). Although the original experiment consisted in duplicate samples every 4 hours during one diel cycle and two circadian cycles, only reads corresponding to the diel cycle were used, corresponding to samples from time-points 12, 16, 20, 24, 28 and 32 (consecutive SRA numbers SRR2452525 to SRR2452536, and SRR2452572 to SRR2452586). Short reads from these 24 libraries were aligned to the tomato genome reference sequence v4.0 using hisat2 v2.1.0 with default parameters (Kim et al., 2015). The number of reads per transcript in the ITAG 4.1 annotation was counted with the featureCounts function using the Rsubread R package with default parameters (Liao et al., 2019). Low expressed transcripts, that contained less than 10 reads across all samples were discarded. The number of reads in the 24330 remaining transcripts was normalized using the vst function in the DEseq2 R package (Love et al., 2014). To improve detection of cycling genes, we split the two replicates from each time-point taken in a single day into two consecutive days by adding 24 hours to the collection time of the second replicate. Cycling genes were identified independently in *S. lycopersicum* and *S. pennellii* using the meta2d function in the R package MetaCycle with parameters minper = 20, maxper = 28, cycMethod = “ARS”, adjustPhase = “predictedPer”, combinePvalue = “fisher”, ARSmle = “auto” and ARSdefaultPer = 24 (Wu et al., 2016). We considered as cycling a total of 5740 genes that presented an adjusted pvalue < 0.05 for the cycling test, an estimated period between 22 and 26 hours both in *S. lycopersicum* and *S. pennellii*, and did not present amplitudes or mean expression values more than 2.5 standard deviations away from the mean (Table S2). Since *S. lycopersicum* presents unusually decelerated rhythms (Müller et al., 2016), cycling genes were assigned the estimated phase from the *S. pennellii* experiment, which was rounded to the nearest integer and subtracted 24 when they exceeded 23 hours.

### Molecular timetable studies

For the molecular timetable representation in Figure 3C, we calculated the average normalized expression of all cycling genes in each genotype, condition and phase bin. For the molecular timetable representation in Figure 3E, we calculated the average log2 fold expression change due to photoperiod of all cycling genes in each genotype and phase bin. Phases of the curves in Figure 3C were calculated using the cosinor function in the psych package in R using normalized expression for all cycling genes in each independent biological replicate (Revelle, 2017). Estimation of significant differences in the response to photoperiod for each genotype were calculated using cosinor phases estimates for each independent replicate and a two way ANOVA in the R package emmeans with genotype and condition as factors (Lenth, 2021).

## Results

### 1. Generation and phenotyping of LNK2+/EID1+ near isogenic lines

In order to investigate how mutations in EID1 and LNK2 interact with each other and how they affect the perception of photoperiod in tomato we generated near isogenic lines (NILs) that contain wild alleles of both genes in a cultivated tomato background. For this we performed several rounds of backcrossing in a set of previously developed NILs containing introgressions from the wild tomato *S. pimpinellifolium* (accession TO937) in a cultivated *S. lycopersicum* (accession Moneymaker) background (Barrantes et al., 2014). We obtained a heterozygous line carrying reduced *S. pimpinellifolium* introgressions at the positions of *EID1* and *LNK2*. Whole genome short read sequencing of this line revealed the presence of 5 introgressions in chromosomes 1, 4, 5 (x2) and 9 (Figure S1). The introgression at chromosome 1 comprised 125 genes and included the *LNK2* locus, while the introgression in chromosome 9 comprised 19 genes, including *EID1*. The progeny of this line was screened for homozygous individuals carrying exclusively the introgression in chromosome 1 (called line LNK2+), the introgression in chromosome 9 (called line EID1+) or both introgressions (EID1+ / LNK2+, called line LE+ hereafter). In addition, all lines selected carried an introgression at the bottom chromosome 4 that includes 208 genes and had been previously detected as a segregation distorter during the generation of the near isogenic line population (Barrantes et al., 2014).

We confirmed the functionality of the *S. pimpinellifolium* alleles of *LNK2* and *EID1* in these lines by characterizing its circadian rhythms in three independent experiments. As expected, wild *LNK2* alleles decreased the long circadian period observed in cultivated tomato but had no significant effect on the phase of leaf movements (Figure 1). Wild *EID1* alleles reduced the circadian phase of cultivated tomato and also had a significant effect reducing its period, albeit not as strongly as *LNK2*. The combination of both wild alleles in the LE+ line was sufficient to restore the period and phase to the values observed in the wild tomato *S. pimpinellifolium*, suggesting that the only mutations altering circadian rhythms in cultivated tomato are the ones present in *LNK2* and *EID1*. We did not observe consistent effects on the amplitude or quality (relative amplitude error) of circadian rhythms (Figure S2). In summary, the set of NILs generated recapitulates the variation in circadian rhythms observed between cultivated tomato and its closest wild relative *S. pimpinellifolium*, and allows studying the interaction between mutations in *EID1* and *LNK2*.

**Figure 1.**
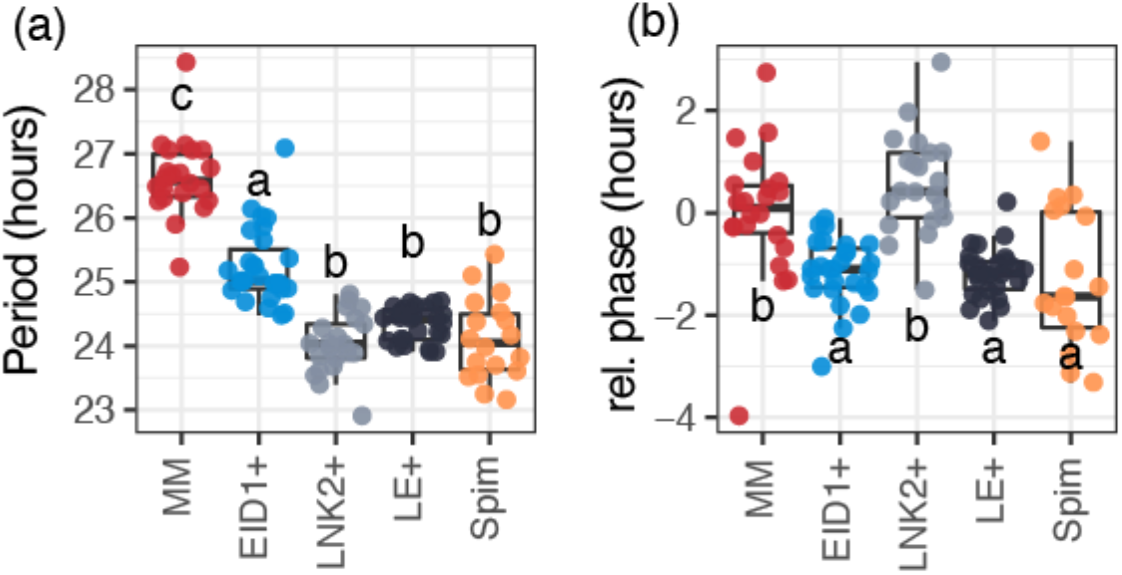
Circadian period **(a)** and phase **(b)** estimates in the near isogenic lines segregating for wild alleles of *EID1* and *LNK2*. Data comes from three independent experiments shown in Figure S2. Relative phase values were obtained by subtracting each phase value from the average of MM in each experiment. MM stands for *S. lycopersicum* cv. MoneyMaker. Spim stands for *S. pimpinellifolium*. Different letters in each boxplot indicate significant differences (p < 0.05, one-way ANOVA and Tukey’s post hoc HSD test).

### 2. Expression analyses

We performed RNA-seq on the near isogenic lines carrying wild alleles of *EID1* and *LNK2* to determine their role in photoperiod perception in tomato. For this, we grew the lines under long day (LD) and short day (SD) conditions, and we collected leaf samples from three biological replicates two hours after lights on (ZT2). This time of the day was chosen to coincide with the highest expression of *EID1* (ZT0) and *LNK2* (ZT4) in a published time course experiment in tomato (Müller et al., 2016)(Figure S3).

We obtained an average of 34.8 million read pairs per sample (minimum of 19.8 million and maximum of 43.8), that were aligned to the tomato reference genome sequence with an average success rate of 92,9%. Principal component analysis (Figure S4) revealed photoperiod as the mayor factor controlling the variance observed among samples (associated to PC1, 65%), while other factors that might be related to the genotype at *LNK2* and *EID1* had smaller effect (associated to PC2, 19%).

### 2.1. Variation in response to genotype

We first investigated if wild alleles of *LNK2* and *EID1* affect gene expression under specific photoperiod by comparing each NIL to *S. lycopersicum* cv Moneymaker (MM) in each photoperiod separately. We found a total of 2316 differentially expressed transcripts (9.95% of the analyzed transcriptome, Figure 2A), with the majority of them (90.75%) showing differences in short days, and only 18.86% showing differences in long days. The largest set of differentially expressed transcripts was found between the EID1+ line and MM in short days (1426 transcripts, Figure 2A), with the majority of those being exclusive to this comparison (1071 transcripts). The second comparison with most abundant differentially expressed transcripts was between the LE+ line and MM in short days (934 transcripts, 480 unique to this comparison). In contrast, the effect of LNK2+ was very limited, affecting only 297 transcripts in short days, suggesting that EID1 has a more important role shaping gene expression than LNK2. Interestingly, only 56 transcripts were differentially expressed between the EID1+ line and MM in LD, indicating that the large effect of EID1 on expression is photoperiod specific.

**Figure 2.**
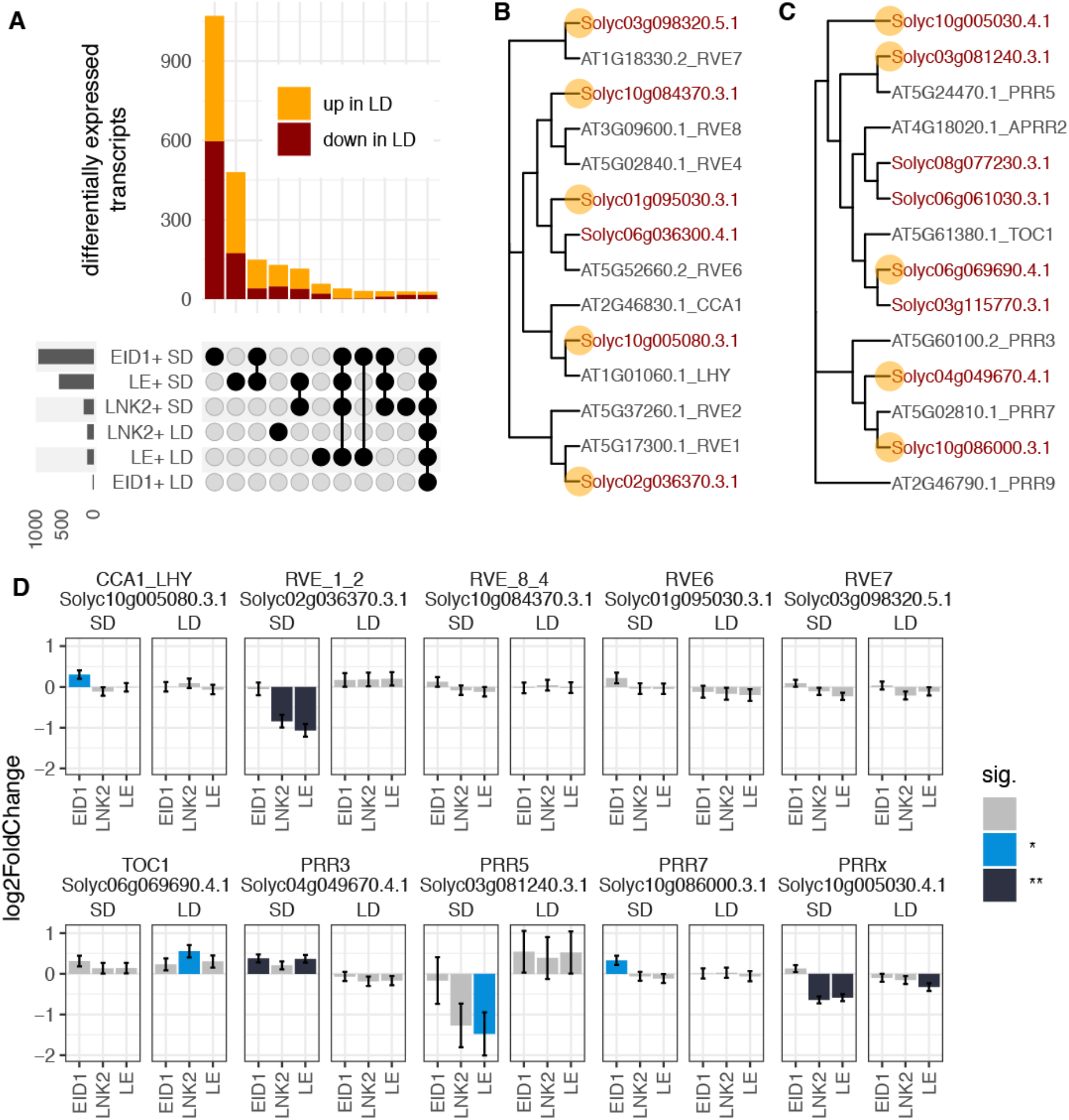
Differentially expressed genes in response to wild alleles of *LNK2* and *EID1*. (A) Bars in the top panel represent the number of differentially expressed genes in each category. Bars in the lower-left panel represent the number of differentially expressed transcripts per genotype and condition. Only sets with more than 20 genes are shown. (B) and (C) Phylogenetic tree with protein sequences from Arabidopsis and tomato for the RVE family (B) and the PRR family (C) of circadian clock genes. Arabidopsis and tomato protein names are highlighted in gray and red respectively. Tomato proteins whose transcripts oscillate during the diel cycle are marked with an orange dot. (D) Log2 fold change in expression (± standard error) between *S. lycopersicum* var MoneyMaker (MM) and each of the near isogenic lines in this work. SD and LD stand for short days and long days. Color of the bars indicate the significance of each log2 fold change, with non-significant values in gray, q<0.05 in blue and q<0.01 in black.

We were surprised by the mild effect of LNK2+ on expression, since LNK proteins in Arabidopsis are involved in transcriptional initiation of clock genes in the PSEUDO RESPONSE REGULATOR (PRR) family such as PRR5 and TOC1, and affects the expression of members of the REVEILLE (RVE) family such as CCA1 or LHY (de Leone et al., 2018; Ma et al., 2018; Pérez-García et al., 2015; Rugnone et al., 2013; Xie et al., 2014). To investigate whether this is the case in tomato, we identified the homologs of the PRR and RVE families from Arabidopsis (Figure 2, panels B and C). While proteins such as RVE7, PRR5 or PRR7 had clear homologs in tomato, some others had been lost (CCA1 / LHY) or had undergone duplication (RVE6 and TOC1). To increase the likelihood of functional homology we considered as homologs only tomato transcripts whose expression oscillated during the diel cycle in a published RNA-seq time-course experiment (Müller et al., 2016).

Among the *RVE* genes in tomato, wild alleles of *LNK2* significantly decreased the expression of the homolog of *RVE1* and *RVE2* in short days, but had no effect on the homolog of *CCA1* and *LHY*, as it does in Arabidopsis (Figure 2). Interestingly, the homolog of *CCA1* and *LHY* in tomato was affected by the addition of wild alleles of *EID1* only in short days. Among the PRRs in tomato, wild alleles of *LNK2* significantly decreased the expression of a *PRRx* transcript for which we could not conclude a closest homolog in Arabidopsis. In concordance with what was observed in Arabidopsis, *LNK2* alleles affected the expression of the homolog of *TOC1*, although this effect was only significant at the 0.05 level in long days, and only in the absence of functional alleles of *EID1*. In addition, wild alleles of *LNK2* strongly decreased the expression of the homolog of *PRR5* in short days, albeit not significantly due to large variation among samples (p=0.16 in LNK2+ and p=0.049 in *LE+*). Finally, wild alleles of *EID1* increased the expression of *PRR3* and *PRR7* in short days. Interestingly, all of the significant differences of expression observed were photoperiod-specific, suggesting a complex seasonal effect in the regulatory function of LNK2. In summary, although some transcripts in the PRR and RVE families in tomato are affected by allelic variation in *LNK2* and *EID1*, we cannot conclude that the mechanism of action of their proteins in tomato are similar to those previously defined in Arabidopsis.

### 2.2. Variation in response to photoperiod

To further explore the interaction of *LNK2* and *EID1* alleles with photoperiod in tomato we analyzed photoperiod sensitivity in each NIL. Fifty percent of the characterized transcriptome (11475 transcripts out of 23269) was significantly affected by photoperiod in at least one genotype (Figure 3). We found roughly the same number of genes upregulated and downregulated by the long day treatment (50.22% upregulated in long days). Moreover, when a gene was differentially expressed in more than one genotype, the direction of the differential expression was consistent between genotypes in 99.5% of the cases.

**Figure 3.**
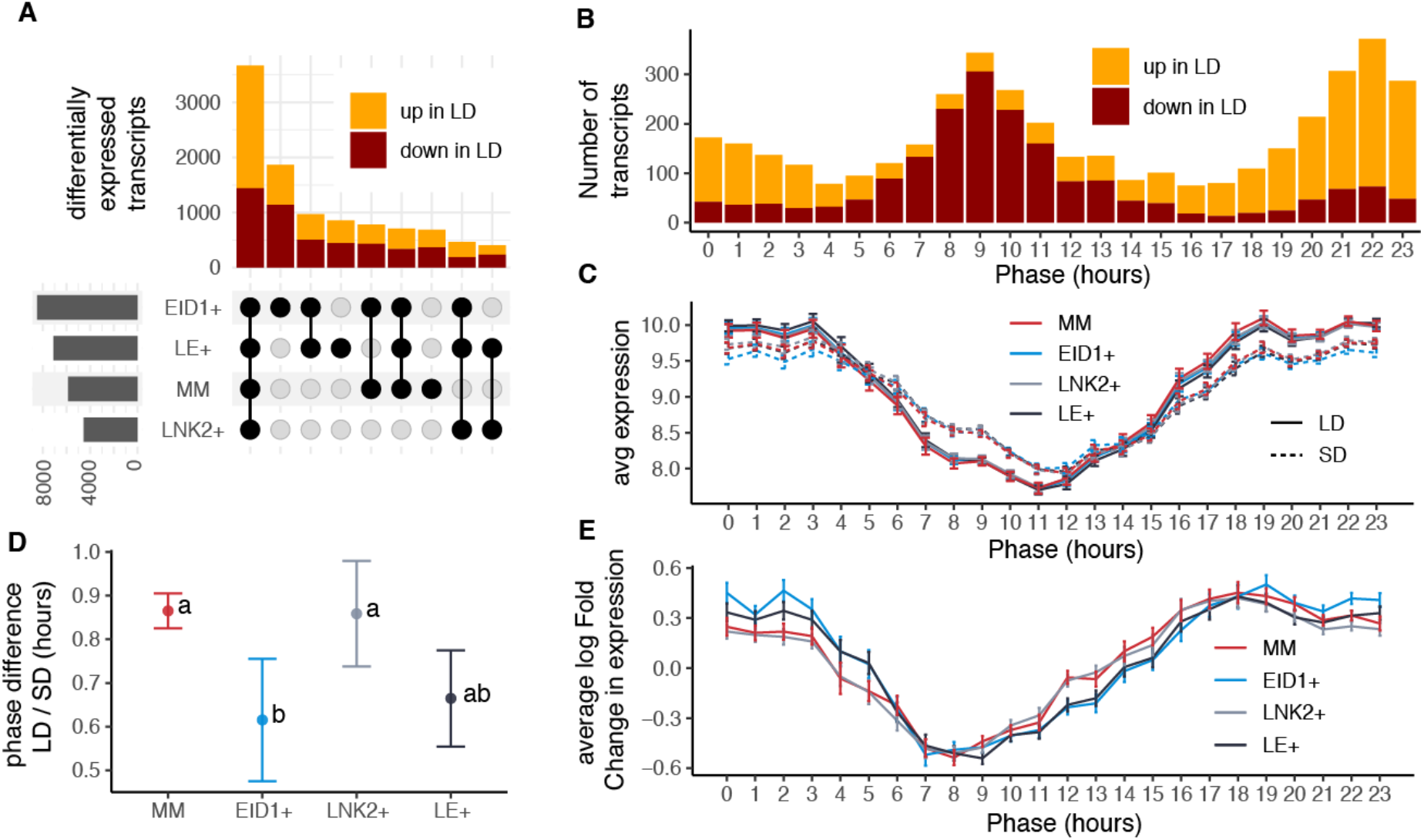
(A) Differential expressed transcripts in response to photoperiod. Bars in the top panel represents the number of differentially expressed genes in each category. Bars in the lower-left panel represent the number of differentially expressed transcripts per genotype. (B) Number of cycling transcripts whose expression is affected by changes in photoperiod grouped by their phase. (C) Average expression of transcripts whose expression oscillated during the day grouped by their phase, genotype and condition. (D) Differences between the estimated phase in long days and short days. Error bars represent the standard deviation of the mean. Different letters in each dot indicate significant differences (p < 0.05, using two-way ANOVA and estimated marginal means). (E) Average log2 fold change in expression induced by photoperiod in cycling genes grouped by genotype and phase of expression. Error bars represent the standard error of the mean.

We divided transcripts significantly responding to photoperiod in groups according to their occurrence in the different genotypes (Figure 3A). The largest group is formed by 3671 transcripts responding to photoperiod in all genotypes, followed by 1866 transcripts affected exclusively in the EID1+ genotype, 978 transcripts deregulated simultaneously in LE+ and EID1+, and 855 differentially expressed in LE+ (Figure 3A). These results suggest that, as observed before and despite a prevalent effect of photoperiod independently of the genotype, the mutation at the *EID1* locus is associated with stronger changes in gene expression, thus stronger photoperiod sensitivity, than the *LNK2* mutation.

Previous studies have determined that changes in photoperiod can impact the expression pattern of transcripts whose expression cycles throughout the day (Dalchau et al., 2010; Michael et al., 2008; Weng et al., 2019). We obtained a list of such transcripts in tomato together with their phase (the time of the day when they show their maximum expression) using previously published RNA-seq data from seedlings from cultivated tomato and its wild relative *S. pennellii* sampled every 4 hours for one day in 12 hours light / 12 hours dark photoperiods (Müller et al., 2016) (Table S2). As observed in other plants, the phases of cycling genes in tomato were not evenly distributed throughout the day, with a majority of transcripts having phases right before dawn (ZT0) or dusk (ZT12) (Figure S5a) (Michael et al., 2008; Weng et al., 2019). Changes in photoperiod significantly affected the expression of more than two thirds of cycling transcripts (4181 out of 6017) whose phases were equally distributed all along the diel cycle (Figure S5b), indicating that changes in photoperiod affect the expression of genes expressed at all times of the day. However, while the majority of cycling genes whose phase occurred at the end of the day were downregulated in long days, cycling genes with phases at the end of the night were mostly upregulated (Figure 3B). Such a marked time of the day specific effect of photoperiod could result from a shift in the phase of circadian rhythms between conditions.

In order to explore this possibility, we evaluated our data using the molecular timetable method that allows phase estimation by visualizing the average expression of cycling genes grouped by their phase (Figure 3C) (Ueda et al., 2004). As expected from samples taken close to dawn (ZT2), cycling transcripts with late night and early morning phases presented higher expression than transcripts whose maximum expression occurs later in the day (Figure 3C). Expression differences between samples in long days and samples in short days were consistent with the results observed before, with long days inducing on average higher expression in dusk genes and lower expression in dawn genes. These results indicate an advanced phase of the circadian clock of tomato under short days compared to long days, as reported for other species before (Dalchau et al., 2010; Michael et al., 2008; Weng et al., 2019). In summary, photoperiod shifts the timing of expression of cycling genes in tomato, thus causing substantial environment specific transcriptional differences.

In order to investigate if wild alleles of *EID1* or *LNK2* affect photoperiodic responses, we estimated the phase differences between the molecular timetable curves in each of our near isogenic lines. Plants in long days have an average of 0.8 hours phase delay with respect to plants grown in short days (Figure 3D). Importantly, phase differences caused by photoperiod were smaller when wild alleles of *EID1* were present, again suggesting that variation in *EID1* but not *LNK2* modulates the phase of cycling genes. We looked for additional confirmation of the effect of photoperiod and *EID1* on the phase of cycling genes by measuring the magnitude of expression changes at each phase bin, since genes with different phases are expected to present larger expression changes at specific times of the day. In Figure 3C we see that the change of phase between long days and short days translates in low expression differences caused by photoperiod around ZT4 and ZT14 and strong differences around ZT8 and ZT20. Representation of the average fold change in expression between photoperiods in each genotype shows that indeed this is the case, with log fold change values at Zt4 and ZT14 approaching 0 (Figure 3E). Moreover, log fold change differences also separate lines carrying different alleles of *EID1* in two groups that present different profiles (Figure 3E). In summary, our results show that photoperiod has a strong effect on the phase of expression of tomato transcripts, and that variation in the *EID1* locus, but not in *LNK2*, modulate this effect.

## Discussion

Here we developed a set of isogenic cultivated tomato lines segregating for wild, functional alleles of *LNK2* and *EID1*, the two genes responsible for delayed circadian rhythms in cultivated tomato (Müller et al., 2016, 2018). Although near isogenic lines containing wild alleles of *EID1* and *LNK2* were developed in our previous works (Müller et al., 2016, 2018), the current lines represent a great improvement over those as a genetic tool to study the role of these genes in tomato physiology. First, the previously existing lines contained large introgressed fragments involving hundreds, if not thousands of genes from the wild species, while the lines developed in this work contain wild alleles in only 125 genes in the region of *LNK2* and 19 genes in the region of *EID1* (Figure S1). It is worth noting that these lines also contain 208 genes with wild alleles at the bottom of chromosome 4, in a region that caused a strong segregation distortion when the population was generated (Barrantes et al., 2014). Despite all our efforts backcrossing these lines, we did not manage to remove this region, although its presence in all lines ensures that it cannot be responsible for the observed differences between lines. Second, the wild donor that we chose to develop the lines in this work is *S. pimpinellifolium*, instead of *S. pennellii* in the previous lines. While *S. pimpinellifolium* is the closest wild relative of cultivated tomato, *S. pennellii* is one of the most distant wild relatives. The closer genetic relationship between donor and acceptor is likely to induce less unintended effects of additional genes present in the introgressions, since they should contain fewer sequence polymorphisms between cultivated and wild alleles.

We confirmed the functionality of the *S. pimpinellifolium* alleles of *EID1* and *LNK2* in the isogenic lines by characterizing their circadian leaf movements (Figure 1). The results coincide with previous work showing that wild alleles of *LNK2* and *EID1* respectively revert the period and phase delay observed in cultivated tomato (Müller et al., 2016, 2018). It is interesting to observe in all three independent experiments showed an additional albeit secondary effect of *EID1* in period (Figure 1). The effect of *EID1* in period coincides in its direction with the shortening of period caused by *LNK2*, confirming that both genes act synergistically and reinforcing the hypothesis of positive selection towards delayed circadian rhythms in cultivated tomato.

The development of these isogenic lines allowed us for the first time to study the interaction of the different alleles of *EID1* and *LNK2*, since no other line was developed before where wild alleles of both genes were present in a cultivated tomato background. The phenotype of the LE+ line resembles perfectly that of the tomato wild relative *S. pimpinellifolium* (Figure S2). This confirms the results of our previously published QTL analysis that identified only two QTLs controlling this trait in a RIL population between cultivated tomato and *S. pimpinellifolium*, predicting that the mutations in *LNK2* and *EID1* are sufficient to generate the delayed circadian rhythm phenotype of cultivated tomato.

We studied the transcriptional changes caused by variation in photoperiod and in the different alleles of *EID1* and *LNK2* at the time when their transcripts are most abundant. Among the two genes studied, *EID1* affected the transcription of many more genes than *LNK2*. The smaller effect of *LNK2* could be explained by redundancy of its protein with other members of its family, that both in Arabidopsis and tomato is composed of 4 members (Figure S6). In Arabidopsis the *lnk2* mutant also shows limited phenotypic differences with the wild type plant as compared with the *lnk1;lnk2* double mutant (Rugnone et al., 2013) or the *lnk1;lnk2;lnk3;lnk4* quadruple mutant (de Leone et al., 2018). In addition, it is also possible that EID1 has a more general role shaping plant expression because of its role regulating photoreceptors (Dieterle et al., 2001; Marrocco et al., 2006), therefore controlling the extensive light signaling network in plants, while LNK2 acts more specifically within the circadian clock and anthocyanin biosynthesis pathways (de Leone et al., 2018; Pérez-García et al., 2015). Nevertheless, we specifically looked for expression variation among the homologs of circadian clock genes in tomato, and did not observe strong differences. One possible explanation for this is that our samples were collected at ZT2, and the changes in expression triggered by *LNK2* are observed later in the day (Ma et al., 2018; Pérez-García et al., 2015; Xie et al., 2014).

The transcriptional response to photoperiod was much larger than the response elicited by the genotype at the *EID1* or *LNK2* loci, affecting roughly half of the tomato transcriptome. This strong response is difficult to compare with previous studies that used other species, methods, significance thresholds and sample collection times. However, scanning of the literature revealed photoperiod effects that range from 50% of the transcriptome in the perennial grass *Panicum hallii* (Weng et al., 2019), to 20 % in Medicago or Arabidopsis (Thomson et al., 2019).

The large number of transcripts affected by photoperiod in our experiment is likely to be caused, at least in part, by the phase change induced by the light treatment. In effect, most transcripts whose expression oscillate during the day are significantly affected by photoperiod following a pattern in which afternoon-expressed genes are downregulated and late-night-expressed genes are upregulated. Using the molecular timetable method we found that this pattern would fit a global phase delay of cycling genes in response to long days, which is similar to what has been observed in other plants such as Arabidopsis (Dalchau et al., 2010; Michael et al., 2008) and *Panicum hallii* (Weng et al., 2019). In contrast, while our method based on a single time point estimates an average phase delay of 0.8 hours (Figure 3D), full time-course datasets performed in Arabidopsis and *Panicum hallii* (Michael et al., 2008; Weng et al., 2019) estimate this phase change to be around 4 hours. It is very likely that this discordance in the estimation of phase differences induced by photoperiod between these studies comes from the low accuracy of our method, which is based in a prediction from a single time point, while the other studies are based on datasets with multiple samples collected at regular intervals during one or various days (Michael et al., 2008; Weng et al., 2019). Phase estimates using full time-course data are more precise due to simpler calculations, but methods that estimate phase from a single time point are gaining popularity because of their low cost and experimental simplification (Braun et al., 2018; Hughey et al., 2016; Ueda et al., 2004). Nevertheless, precise photoperiod-induced phase change estimation in tomato would require measuring expression from samples collected at regular intervals during at least one-day in both conditions.

In summary, our work reveals a role of EID1 but not LNK2 in the perception of photoperiod in tomato through modulation of the phase of expression of cycling genes. Changes in photoperiod sensibility could have been important for tomato to better adapt to variation in day length in latitudes outside the tropics.

## Data availability

Sequence data generated for this manuscript is available at the Short Read Archive (https://www.ncbi.nlm.nih.gov/sra), under project PRJNA797239. All other relevant data can be found within the manuscript and its supporting materials.

## Acknowledgments

We thank Antonio Monforte for sharing the *S. pimpinellifolium* introgression line population used to construct the near isogenic lines. We thank Michel Le-Brusq, Patrick Grillot, Michel Burtin and Lilian Dahuron for help growing the plants. We Thank Nazneen Badroudine, Maria Jesus Lacruz, Adrien Léonardi, Magali Nawrocki-Serin, Vanessa Pacé-Faria, Stéphane Raude and Stéphanie Zimmermann for help with administrative tasks. We thank Niels A. Müller and Paloma Mas for discussion and critical reading of the manuscript.

## Supplementary Figures

**Figure S1.**
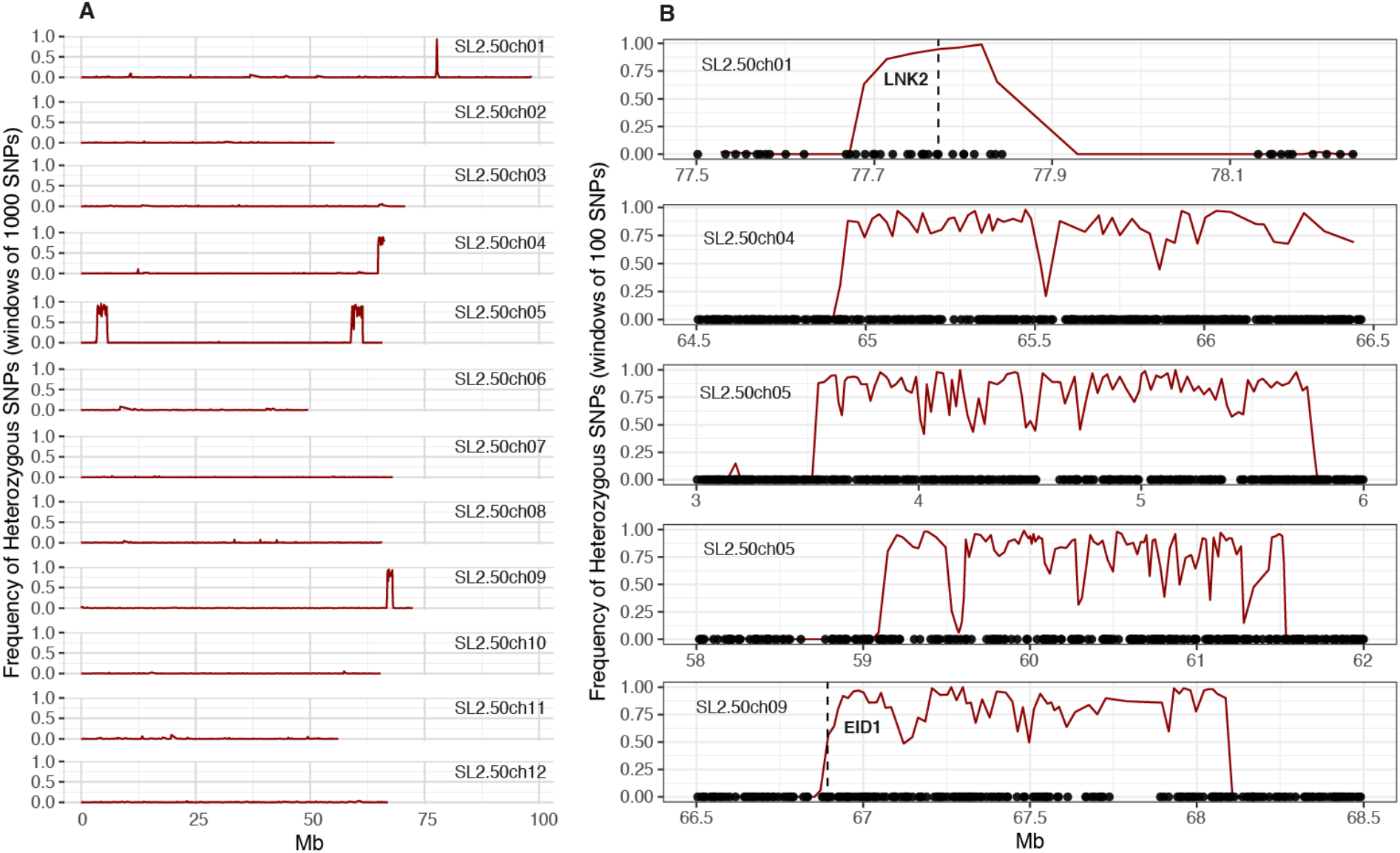
Whole genome genotyping of a heterozygous line containing reduced introgressions at the positions of *LNK2* and *EID1*. (A) The red line indicates the frequency of heterozygous SNPs in 1000-SNP windows along the 12 tomato chromosomes. (B) Zoom-in view in the chromosomal regions where the heterozygous line presented introgressions from *S. pimpinellifolium*. Red lines indicate the frequency of heterozygous SNPs in 100-SNP windows. The location of *LNK2* and *EID1* is indicated with a vertical line. The location of genes in each region is indicated with black dots along the x axis.

**Figure S2.**
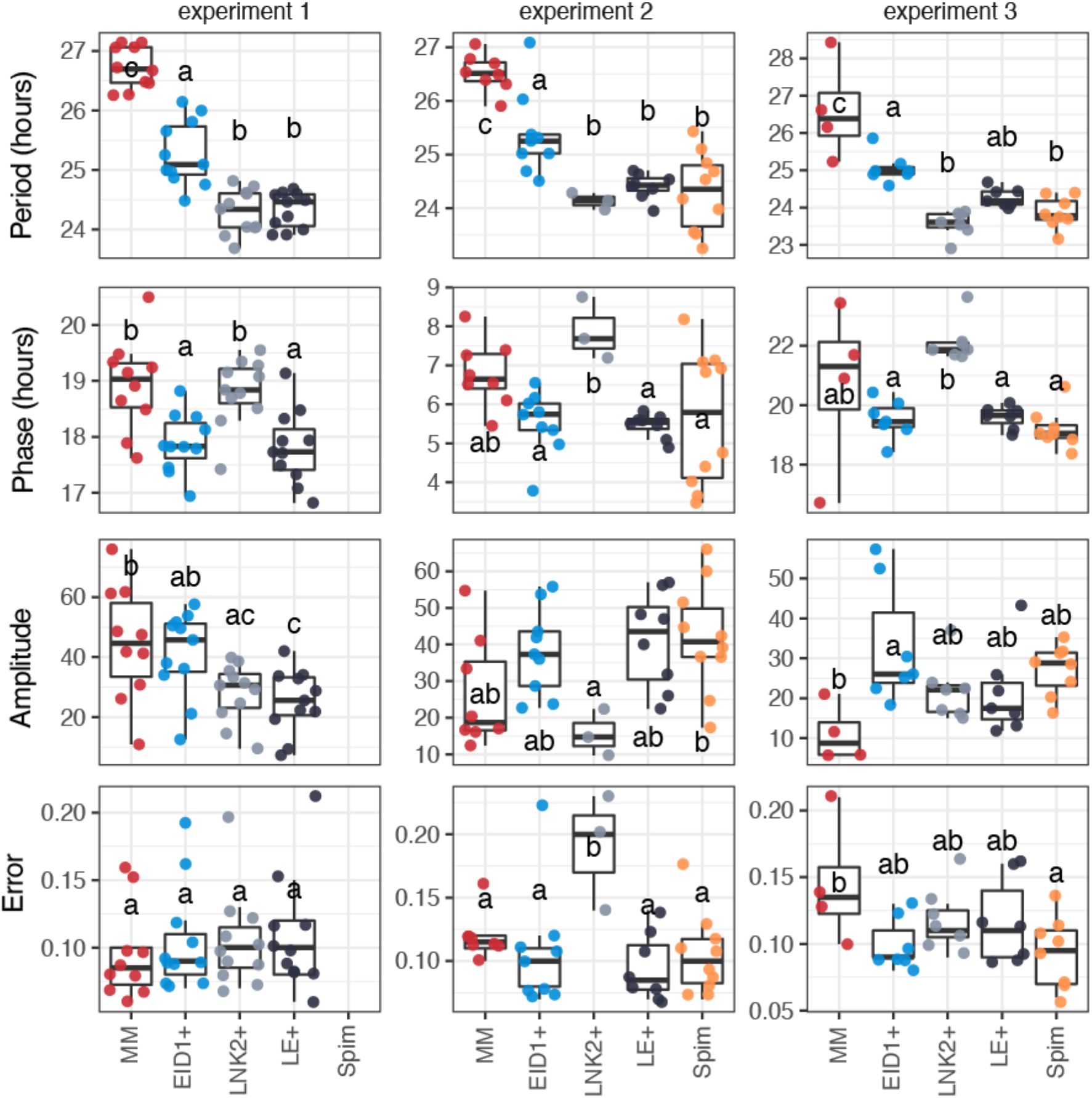
Circadian parameters in the near isogenic lines generated. Three independent experiments are shown in columns and Period, Phase, Amplitude and Relative Amplitude Error in rows. The wild species *S. pimpinellifolium* was not included in the first experiment. Different letters in each boxplot indicate significant differences (P < 0.05, one-way ANOVA and Tukey’s post hoc HSD test).

**Figure S3.**
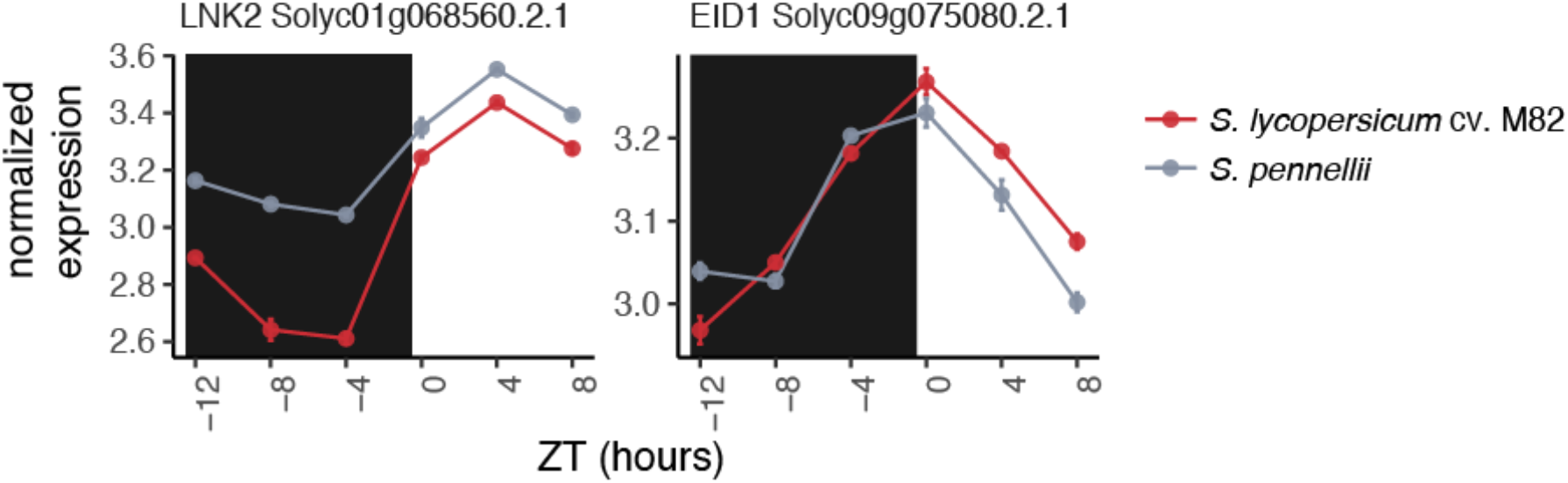
Expression oscillation of *LNK2* and *EID1* in tomato. Data was obtained from RNA-seq data published in Müller et al 2016. Plants were grown in 12:12 light/dark and 20:18 °C temperature cycles and leaf samples collected from 7-day old seedlings every 4 hours. Read counts on each gene are normalized by gene length and sample size.

**Figure S4.**
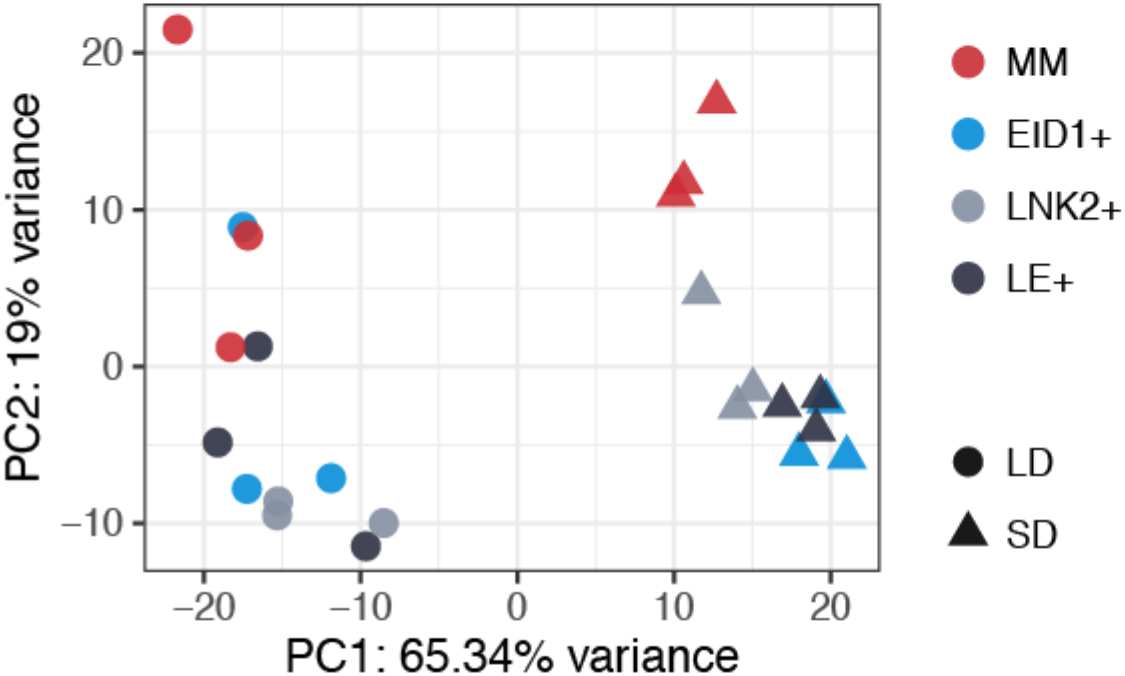
Principal component analysis of expression values from the RNA-seq experiment in the near isogenic lines segregating for wild alleles of *EID1* and *LNK2*. Only transcripts with more than 10 reads across all samples were included in the analysis.

**Figure S5.**
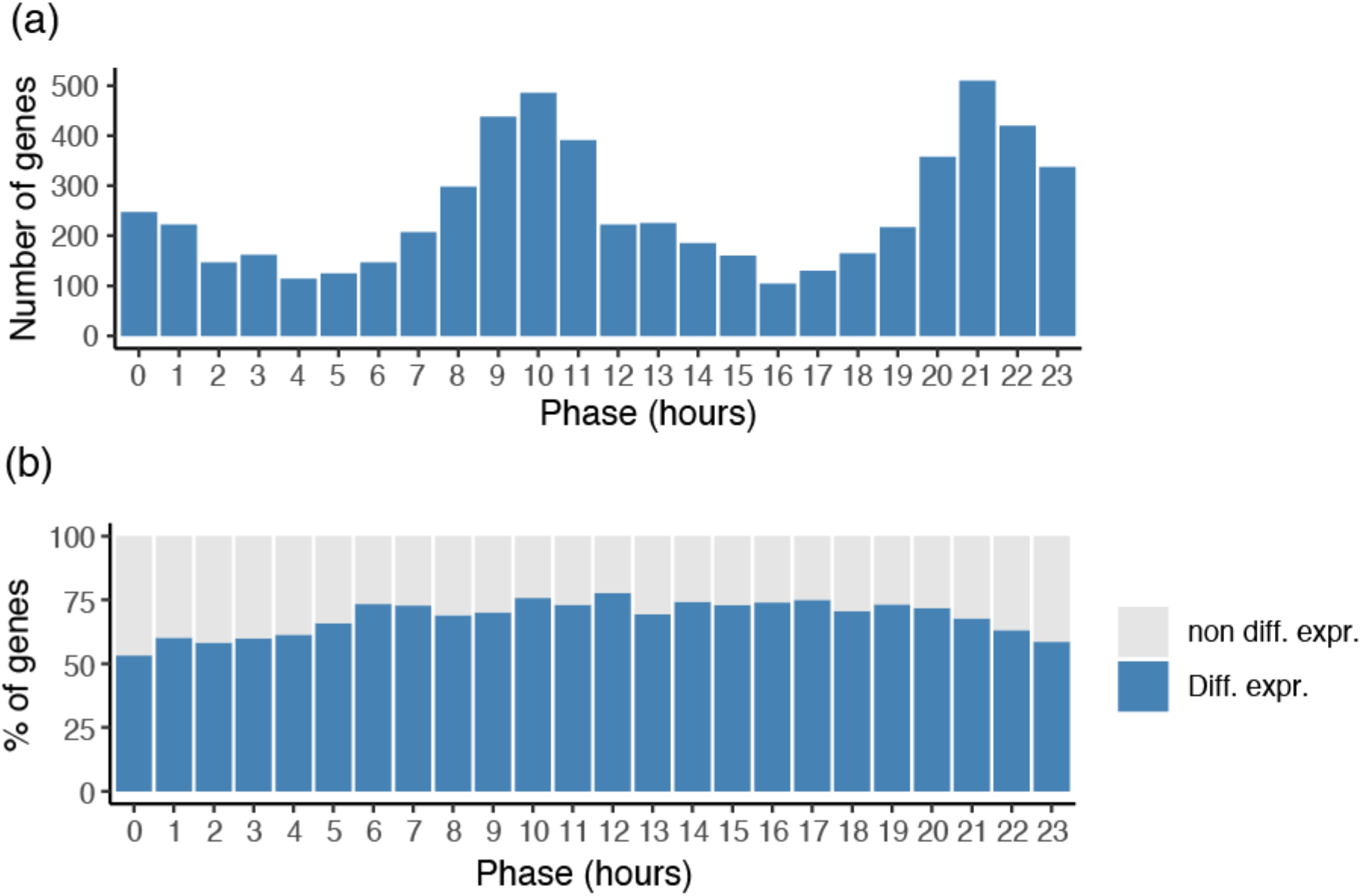
Phase and percent of differentially expressed genes among cycling genes in tomato. (a) Phase distribution of the 6017 transcripts whose expression oscillates during the diel cycle in *S. lycopersicum* and *S. pennellii*. (b) Percentage of transcripts in (a) whose expression was significantly altered by photoperiod in our experiment.

**Figure S6.**
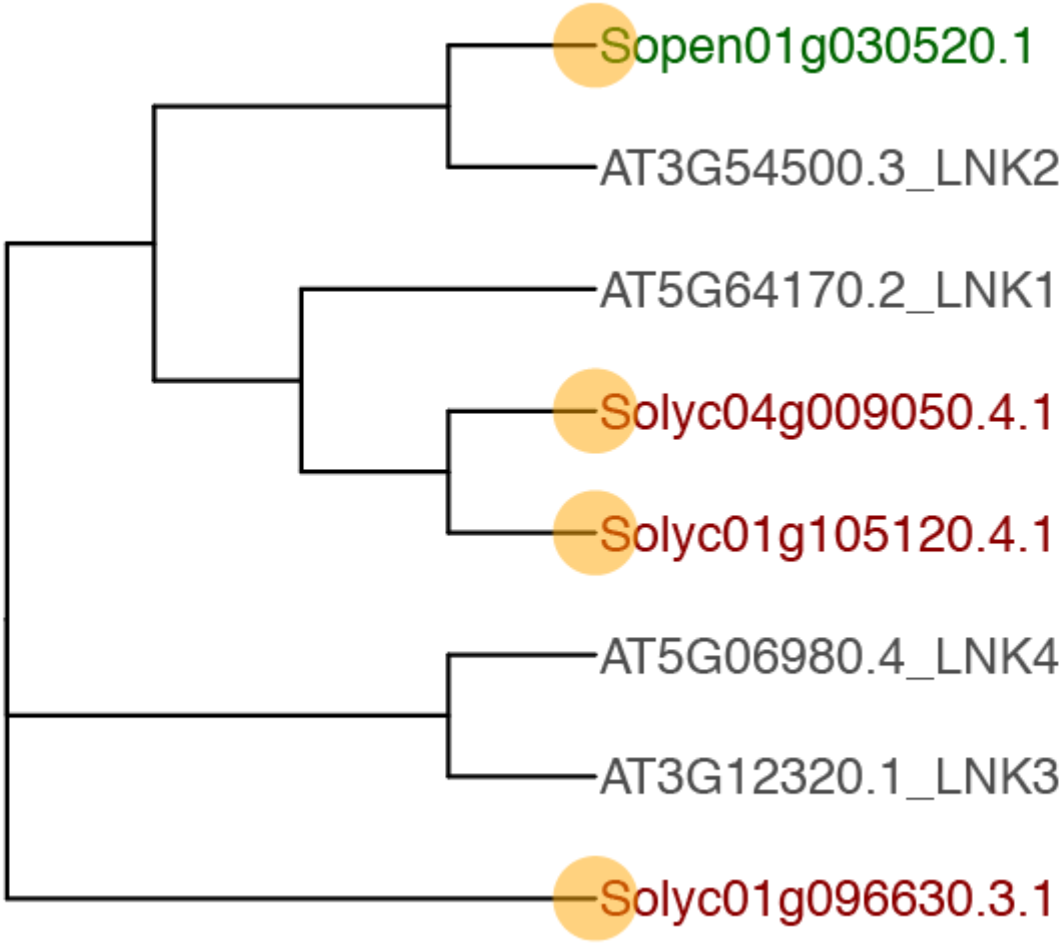
Phylogenetic tree from protein sequence alignments for genes belonging to the LNK family in tomato and Arabidopsis. For *LNK2*, the sequence from the wild tomato species *S. pennellii* is used because of the large deletion present in this gene in cultivated tomato. Arabidopsis, cultivated tomato and *S. pennellii* protein names are highlighted in gray, red and green respectively. Tomato proteins whose transcript oscillates during the diel cycle are marked with an orange dot.

## Notes

### Competing Interest Statement

The authors have declared no competing interest.

## References

Aflitos, S., Schijlen, E., de Jong, H., de Ridder, D., Smit, S., Finkers, R., Wang, J., Zhang, G., Li, N., Mao, L., et al. (2014). Exploring genetic variation in the tomato (Solanum section Lycopersicon) clade by whole-genome sequencing. Plant J. Cell Mol. Biol. 80, 136–148.

Barrantes, W., Fernández-del-Carmen, A., López-Casado, G., González-Sánchez, M.Á., Fernández-Muñoz, R., Granell, A., and Monforte, A.J. (2014). Highly efficient genomics-assisted development of a library of introgression lines of Solanum pimpinellifolium. Mol. Breed. 34, 1817–1831.

Barrantes, W., López-Casado, G., García-Martínez, S., Alonso, A., Rubio, F., Ruiz, J.J., Fernández-Muñoz, R., Granell, A., and Monforte, A.J. (2016). Exploring New Alleles Involved in Tomato Fruit Quality in an Introgression Line Library of Solanum pimpinellifolium. Front. Plant Sci. 7, 1172.

Bendix, C., Marshall, C.M., and Harmon, F.G. (2015). Circadian Clock Genes Universally Control Key Agricultural Traits. Mol. Plant 8, 1135–1152.

Braun, R., Kath, W.L., Iwanaszko, M., Kula-Eversole, E., Abbott, S.M., Reid, K.J., Zee, P.C., and Allada, R. (2018). Universal method for robust detection of circadian state from gene expression. Proc. Natl. Acad. Sci. 115, E9247–E9256.

Camacho, C., Coulouris, G., Avagyan, V., Ma, N., Papadopoulos, J., Bealer, K., and Madden, T.L. (2009). BLAST+: architecture and applications. BMC Bioinformatics 10, 421.

Dalchau, N., Hubbard, K.E., Robertson, F.C., Hotta, C.T., Briggs, H.M., Stan, G.-B., Gonçalves, J.M., and Webb, A.A.R. (2010). Correct biological timing in Arabidopsis requires multiple light-signaling pathways. Proc. Natl. Acad. Sci. U. S. A. 107, 13171–13176.

Dieterle, M., Zhou, Y.C., Schäfer, E., Funk, M., and Kretsch, T. (2001). EID1, an F-box protein involved in phytochrome A-specific light signaling. Genes Dev. 15, 939–944.

Edgar, R.S., Green, E.W., Zhao, Y., van Ooijen, G., Olmedo, M., Qin, X., Xu, Y., Pan, M., Valekunja, U.K., Feeney, K.A., et al. (2012). Peroxiredoxins are conserved markers of circadian rhythms. Nature 485, 459–464.

Faure, S., Turner, A.S., Gruszka, D., Christodoulou, V., Davis, S.J., von Korff, M., and Laurie, D.A. (2012). Mutation at the circadian clock gene EARLY MATURITY 8 adapts domesticated barley (Hordeum vulgare) to short growing seasons. Proc. Natl. Acad. Sci. U. S. A. 109, 8328–8333.

Greenham, K., Lou, P., Puzey, J.R., Kumar, G., Arnevik, C., Farid, H., Willis, J.H., and McClung, C.R. (2017). Geographic Variation of Plant Circadian Clock Function in Natural and Agricultural Settings. J. Biol. Rhythms 32, 26–34.

Harmer, S.L. (2009). The circadian system in higher plants. Annu. Rev. Plant Biol. 60, 357–377.

Hughey, J.J., Hastie, T., and Butte, A.J. (2016). ZeitZeiger: supervised learning for high-dimensional data from an oscillatory system. Nucleic Acids Res. 44, e80.

Hut, R.A., Paolucci, S., Dor, R., Kyriacou, C.P., and Daan, S. (2013). Latitudinal clines: an evolutionary view on biological rhythms†,‡. Proc. R. Soc. B Biol. Sci. 280, 20130433.

Jones, M.A. (2009). Entrainment of the Arabidopsis Circadian Clock. J. Plant Biol. 52, 202–209.

Kim, D., Langmead, B., and Salzberg, S.L. (2015). HISAT: a fast spliced aligner with low memory requirements. Nat. Methods 12, 357–360.

Langmead, B., and Salzberg, S.L. (2012). Fast gapped-read alignment with Bowtie 2. Nat. Methods 9, 357–359.

Lenth, R.V. (2021). emmeans: Estimated Marginal Means, aka Least-Squares Means.

de Leone, M.J., Hernando, C.E., Romanowski, A., García-Hourquet, M., Careno, D., Casal, J., Rugnone, M., Mora-García, S., and Yanovsky, M.J. (2018). The LNK Gene Family: At the Crossroad between Light Signaling and the Circadian Clock. Genes 10.

Li, H., Handsaker, B., Wysoker, A., Fennell, T., Ruan, J., Homer, N., Marth, G., Abecasis, G., Durbin, R., and 1000 Genome Project Data Processing Subgroup (2009). The Sequence Alignment/Map format and SAMtools. Bioinforma. Oxf. Engl. 25, 2078–2079.

Li, M.-W., Liu, W., Lam, H.-M., and Gendron, J.M. (2019). Characterization of Two Growth Period QTLs Reveals Modification of PRR3 Genes During Soybean Domestication. Plant Cell Physiol. 60, 407–420.

Liao, Y., Smyth, G.K., and Shi, W. (2019). The R package Rsubread is easier, faster, cheaper and better for alignment and quantification of RNA sequencing reads. Nucleic Acids Res. 47, e47–e47.

Lin, T., Zhu, G., Zhang, J., Xu, X., Yu, Q., Zheng, Z., Zhang, Z., Lun, Y., Li, S., Wang, X., et al. (2014). Genomic analyses provide insights into the history of tomato breeding. Nat. Genet. 46, 1220–1226.

Love, M.I., Huber, W., and Anders, S. (2014). Moderated estimation of fold change and dispersion for RNA-seq data with DESeq2. Genome Biol. 15, 550.

Lu, S., Zhao, X., Hu, Y., Liu, S., Nan, H., Li, X., Fang, C., Cao, D., Shi, X., Kong, L., et al. (2017). Natural variation at the soybean J locus improves adaptation to the tropics and enhances yield. Nat. Genet. 49, 773–779.

Ma, Y., Gil, S., Grasser, K.D., and Mas, P. (2018). Targeted Recruitment of the Basal Transcriptional Machinery by LNK Clock Components Controls the Circadian Rhythms of Nascent RNAs in Arabidopsis. Plant Cell 30, 907–924.

Marrocco, K., Zhou, Y., Bury, E., Dieterle, M., Funk, M., Genschik, P., Krenz, M., Stolpe, T., and Kretsch, T. (2006). Functional analysis of EID1, an F-box protein involved in phytochrome A-dependent light signal transduction. Plant J. Cell Mol. Biol. 45, 423–438.

McClung, C.R. (2021). Circadian Clock Components Offer Targets for Crop Domestication and Improvement. Genes 12, 374.

McKenna, A., Hanna, M., Banks, E., Sivachenko, A., Cibulskis, K., Kernytsky, A., Garimella, K., Altshuler, D., Gabriel, S., Daly, M., et al. (2010). The Genome Analysis Toolkit: a MapReduce framework for analyzing next-generation DNA sequencing data. Genome Res. 20, 1297–1303.

Michael, T.P., Salomé, P.A., Yu, H.J., Spencer, T.R., Sharp, E.L., McPeek, M.A., Alonso, J.M., Ecker, J.R., and McClung, C.R. (2003). Enhanced fitness conferred by naturally occurring variation in the circadian clock. Science 302, 1049–1053.

Michael, T.P., Mockler, T.C., Breton, G., McEntee, C., Byer, A., Trout, J.D., Hazen, S.P., Shen, R., Priest, H.D., Sullivan, C.M., et al. (2008). Network discovery pipeline elucidates conserved time-of-day-specific cis-regulatory modules. PLoS Genet. 4, e14.

Miwa, K., Serikawa, M., Suzuki, S., Kondo, T., and Oyama, T. (2006). Conserved expression profiles of circadian clock-related genes in two Lemna species showing long-day and short-day photoperiodic flowering responses. Plant Cell Physiol. 47, 601–612.

Müller, N.A., and Jiménez-Gómez, J.M. (2016). Analysis of Circadian Leaf Movements. Methods Mol. Biol. Clifton NJ 1398, 71–79.

Müller, N.A., Wijnen, C.L., Srinivasan, A., Ryngajllo, M., Ofner, I., Lin, T., Ranjan, A., West, D., Maloof, J.N., Sinha, N.R., et al. (2016). Domestication selected for deceleration of the circadian clock in cultivated tomato. Nat. Genet. 48, 89–93.

Müller, N.A., Zhang, L., Koornneef, M., and Jiménez-Gómez, J.M. (2018). Mutations in EID1 and LNK2 caused light-conditional clock deceleration during tomato domestication. Proc. Natl. Acad. Sci. U. S. A.

Nakamichi, N., Kiba, T., Henriques, R., Mizuno, T., Chua, N.-H., and Sakakibara, H. (2010). PSEUDO-RESPONSE REGULATORS 9, 7, and 5 are transcriptional repressors in the Arabidopsis circadian clock. Plant Cell 22, 594–605.

Paradis, E., and Schliep, K. (2019). ape 5.0: an environment for modern phylogenetics and evolutionary analyses in R. Bioinforma. Oxf. Engl. 35, 526–528.

Pérez-García, P., Ma, Y., Yanovsky, M.J., and Mas, P. (2015). Time-dependent sequestration of RVE8 by LNK proteins shapes the diurnal oscillation of anthocyanin biosynthesis. Proc. Natl. Acad. Sci. U. S. A. 112, 5249–5253.

Pin, P.A., Zhang, W., Vogt, S.H., Dally, N., Büttner, B., Schulze-Buxloh, G., Jelly, N.S., Chia, T.Y.P., Mutasa-Göttgens, E.S., Dohm, J.C., et al. (2012). The role of a pseudo-response regulator gene in life cycle adaptation and domestication of beet. Curr. Biol. CB 22, 1095–1101.

Rawat, R., Takahashi, N., Hsu, P.Y., Jones, M.A., Schwartz, J., Salemi, M.R., Phinney, B.S., and Harmer, S.L. (2011). REVEILLE8 and PSEUDO-REPONSE REGULATOR5 form a negative feedback loop within the Arabidopsis circadian clock. PLoS Genet. 7, e1001350.

Revelle, W. (2017). psych: Procedures for Personality and Psychological Research.

Roden, L.C., Song, H.-R., Jackson, S., Morris, K., and Carre, I.A. (2002). Floral responses to photoperiod are correlated with the timing of rhythmic expression relative to dawn and dusk in Arabidopsis. Proc. Natl. Acad. Sci. U. S. A. 99, 13313–13318.

Rugnone, M.L., Faigón Soverna, A., Sanchez, S.E., Schlaen, R.G., Hernando, C.E., Seymour, D.K., Mancini, E., Chernomoretz, A., Weigel, D., Más, P., et al. (2013). LNK genes integrate light and clock signaling networks at the core of the Arabidopsis oscillator. Proc. Natl. Acad. Sci. U. S. A. 110, 12120–12125.

Sanchez, S.E., Rugnone, M.L., and Kay, S.A. (2020). Light Perception: A Matter of Time. Mol. Plant 13, 363–385.

Schneider, C.A., Rasband, W.S., and Eliceiri, K.W. (2012). NIH Image to ImageJ: 25 years of image analysis. Nat. Methods 9, 671–675.

Steed, G., Ramirez, D.C., Hannah, M.A., and Webb, A.A.R. (2021). Chronoculture, harnessing the circadian clock to improve crop yield and sustainability. Science 372, eabc9141.

Thomson, G., Taylor, J., and Putterill, J. (2019). The transcriptomic response to a short day to long day shift in leaves of the reference legume Medicago truncatula. PeerJ 7, e6626.

Turner, A., Beales, J., Faure, S., Dunford, R.P., and Laurie, D.A. (2005). The pseudo-response regulator Ppd-H1 provides adaptation to photoperiod in barley. Science 310, 1031–1034.

Ueda, H.R., Chen, W., Minami, Y., Honma, S., Honma, K., Iino, M., and Hashimoto, S. (2004). Molecular-timetable methods for detection of body time and rhythm disorders from single-time-point genome-wide expression profiles. Proc. Natl. Acad. Sci. 101, 11227–11232.

Watanabe, S., Xia, Z., Hideshima, R., Tsubokura, Y., Sato, S., Yamanaka, N., Takahashi, R., Anai, T., Tabata, S., Kitamura, K., et al. (2011). A map-based cloning strategy employing a residual heterozygous line reveals that the GIGANTEA gene is involved in soybean maturity and flowering. Genetics 188, 395–407.

Webb, A.A.R., Seki, M., Satake, A., and Caldana, C. (2019). Continuous dynamic adjustment of the plant circadian oscillator. Nat. Commun. 10, 550.

Weller, J.L., Liew, L.C., Hecht, V.F.G., Rajandran, V., Laurie, R.E., Ridge, S., Wenden, B., Vander Schoor, J.K., Jaminon, O., Blassiau, C., et al. (2012). A conserved molecular basis for photoperiod adaptation in two temperate legumes. Proc. Natl. Acad. Sci. U. S. A. 109, 21158–21163.

Weng, X., Lovell, J.T., Schwartz, S.L., Cheng, C., Haque, T., Zhang, L., Razzaque, S., and Juenger, T.E. (2019). Complex interactions between day length and diurnal patterns of gene expression drive photoperiodic responses in a perennial C4 grass. Plant Cell Environ. 42, 2165–2182.

Wu, G., Anafi, R.C., Hughes, M.E., Kornacker, K., and Hogenesch, J.B. (2016). MetaCycle: an integrated R package to evaluate periodicity in large scale data. Bioinformatics 32, 3351–3353.

Xie, Q., Wang, P., Liu, X., Yuan, L., Wang, L., Zhang, C., Li, Y., Xing, H., Zhi, L., Yue, Z., et al. (2014). LNK1 and LNK2 are transcriptional coactivators in the Arabidopsis circadian oscillator. Plant Cell 26, 2843–2857.

Yu, G., Smith, D.K., Zhu, H., Guan, Y., and Lam, T.T.-Y. (2016). ggtree: an r package for visualization and annotation of phylogenetic trees with their covariates and other associated data. Methods Ecol. Evol. 8, 28–36.

Zielinski, T., Moore, A.M., Troup, E., Halliday, K.J., and Millar, A.J. (2014). Strengths and limitations of period estimation methods for circadian data. PloS One 9, e96462.

